# *APOE4* and *INPP5D* converge on membrane mechanics to regulate endocytosis in human astrocytes

**DOI:** 10.64898/2026.05.22.726347

**Authors:** Artur S. Gevorgyan, Lila M. Gordon, Christian A. Combs, Adrian Lita, Linda G. Yang, Carla Cuní-López, Jessica T. Root, Mioara Larion, Justin W. Taraska, Priyanka S. Narayan

## Abstract

Disrupted endocytosis is an early feature of Alzheimer’s disease (AD), but how genetic risk factors functionally impact this pathway remains unclear. Using isogenic human iPSC-derived astrocytes, we show that the AD risk variant, *APOE4,* impairs clathrin-mediated membrane curvature, clathrin-mediated endocytosis, and alters plasma membrane lipid saturation and tension. Compared to *APOE3* astrocytes, *APOE4* cells display an accumulation of flat clathrin structures, reduced maturation of clathrin-coated pits, decreased early endosomes, and reduced endocytic uptake. We then identify the AD risk gene *INPP5D* as a modifier that restores early endocytosis in *APOE4* astrocytes by promoting clathrin curvature and maturation through a mechanism distinct from membrane tension regulation. Beyond effects on trafficking, *INPP5D* overexpression also reduces lipid droplet accumulation and attenuates inflammatory signaling, linking membrane dynamics to disease-associated astrocytic phenotypes. Together, these findings establish altered membrane mechanics as a proximal consequence of the *APOE4* variant and identify endocytosis as a common node linking AD risk genes.

## Introduction

Disrupted endocytosis is an early pathological hallmark of Alzheimer’s disease (AD)^1,2^. Genome-wide association studies (GWAS) for late-onset AD, have identified many single nucleotide polymorphisms (SNPs) in or near endocytosis-related genes including, *BIN1, INPP5D, PICALM, CD2AP*, among others^1,3^. Studies of postmortem human brain AD pathology and rodent AD models have identified endosomal dysfunction as an early pathology of disease even preceding amyloid deposition^4–11^. Moreover, the strongest genetic risk factor for late-onset AD, *APOE4,* has also been linked with disrupted endocytosis in mouse and human cellular systems^7,12,13^.

*APOE* encodes apolipoprotein E, a lipid transport protein predominantly produced by astrocytes in the central nervous system^14,15^. The two most common alleles of *APOE* are *APOE3* (neutral with respect to AD risk) and *APOE4* (C112R), which increases AD risk by up to 15-fold in homozygous individuals^16^. Previous work identified that *APOE4* induces a defect in clathrin-mediated endocytosis (CME) in astrocytes^12^. CME is central to intercellular communication and nutrient uptake in many cell types. Astrocytes use CME to receive signals from multiple cell types throughout the central nervous system (CNS), and disrupted endocytosis impacts both cell autonomous and non-autonomous processes. Although the *APOE4* genotype has been linked to disrupted endocytosis for years, the molecular underpinnings of this connection are still not understood^4^. Since *APOE* encodes a highly abundant brain lipid transport protein, mainly produced and secreted by astrocytes in CNS, we hypothesized that *APOE4* could lead to endocytic disruptions in astrocytes through *APOE4-*associated changes to membrane lipid compositions and consequently membrane dynamics.

In this work, we discovered that human induced pluripotent stem cell (iPSC)-derived astrocytes harboring the *APOE4* genotype display altered clathrin structures at the plasma membrane which link to functional defects throughout the endocytic pathway. We also show that *APOE4* impacts plasma membrane lipid composition and tension which could lead to impaired endocytosis. Additionally, we found that increasing *INPP5D* expression alleviates the functional defects of *APOE4* on plasma membrane clathrin and consequently, endocytosis in astrocytes. *INPP5D* encodes SHIP1, a phosphatidylinositol phosphatase that has previously been shown to be a negative regulator of TREM2 activation, phagocytosis, disrupted autophagy, and inflammasome activation in microglia^17–20^. Together our work shows consequences of *APOE4* expression on astrocyte plasma membrane lipid composition and organelle function in the context of AD risk and centers endocytosis as a convergent node linking multiple AD risk genes.

## Results

All experiments were conducted in astrocytes derived from two isogenic sets of human iPSC lines, KOLF2.1J^21^ and AG09173^7^. Each set contains an *APOE3* homozygous parental line and a CRISPR-Cas9-generated *APOE4* homozygous isogenic counterpart. All iPSC lines used were karyotypically normal^22,23^. Once differentiated to astrocytes using small molecule-guided protocols^7^, cells from both isogenic sets expressed the canonical astrocyte markers: S100b and GFAP (Figure S1A). All characterizations were performed in media conditions without serum, as serum exposure has been found to alter several key astrocyte functions^24–26^.

### *APOE4* decreases clathrin curvature generation at the plasma membrane

Endocytic dysfunction is an early feature of Alzheimer’s disease pathology and is influenced by multiple AD risk genes, including *APOE4*^4,6,7,12^. Although previous work has shown that *APOE4* impairs CME in astrocytes^12^, the molecular mechanisms of this defect remain unclear. Clathrin-mediated endocytosis depends on the nucleation and maturation of clathrin-coated pits (CCPs) at the plasma membrane. We first examined whether *APOE4* disrupts the assembly and morphology of early CCPs in astrocytes. Platinum replica electron microscopy (PREM)^27^ allows for the detailed visualization and characterization of clathrin lattice structures on the inner leaflet of the plasma membrane^28^. Using PREM, we characterized the morphology and distribution of clathrin on the cytoplasmic plasma membrane (PM) of *APOE3* and *APOE4* astrocytes (Figure 1A). We observed a variety of flat, domed, and spherical morphologies of clathrin in both *APOE3* and *APOE4* astrocytes, which correspond to the various stages of clathrin-coated pit maturation (Figure 1B). Using unbiased AI-based identification, we segmented all clathrin structures into three general structural categories of flat, domed, and spherical clathrin (Figure 1C). *APOE4* astrocyte membranes had a greater number of total clathrin structures (Figure 1D) than *APOE3* astrocytes. Of these structures, there were greater numbers of flat clathrin structures in *APOE4* astrocytes compared to *APOE3* astrocyte membranes (Figure 1E), but no changes in flat clathrin size were noted between *APOE3* and *APOE4* astrocyte membranes (Figure 1F). There were no statistically significant differences in the number or sizes of domed or spherical clathrin found between *APOE3* and *APOE4* astrocytes (Figure 1G-J). We also observed no clear differences in the percentage of clathrin area per cell area between *APOE3* and *APOE4* astrocytes (Figure S1B-E). One possible model of CME relies on the ordered maturation of clathrin lattices from flat to domed to spherical vesicles^28^. Therefore, we quantified the efficiency of these transitions using the ratio of domes:flats and spheres:domes. Compared to *APOE3* astrocytes, *APOE4* astrocytes showed reduced conversion of flat to domed clathrin (Figure 1K) and domed to spherical clathrin (Figure 1L).

**Figure 1:**
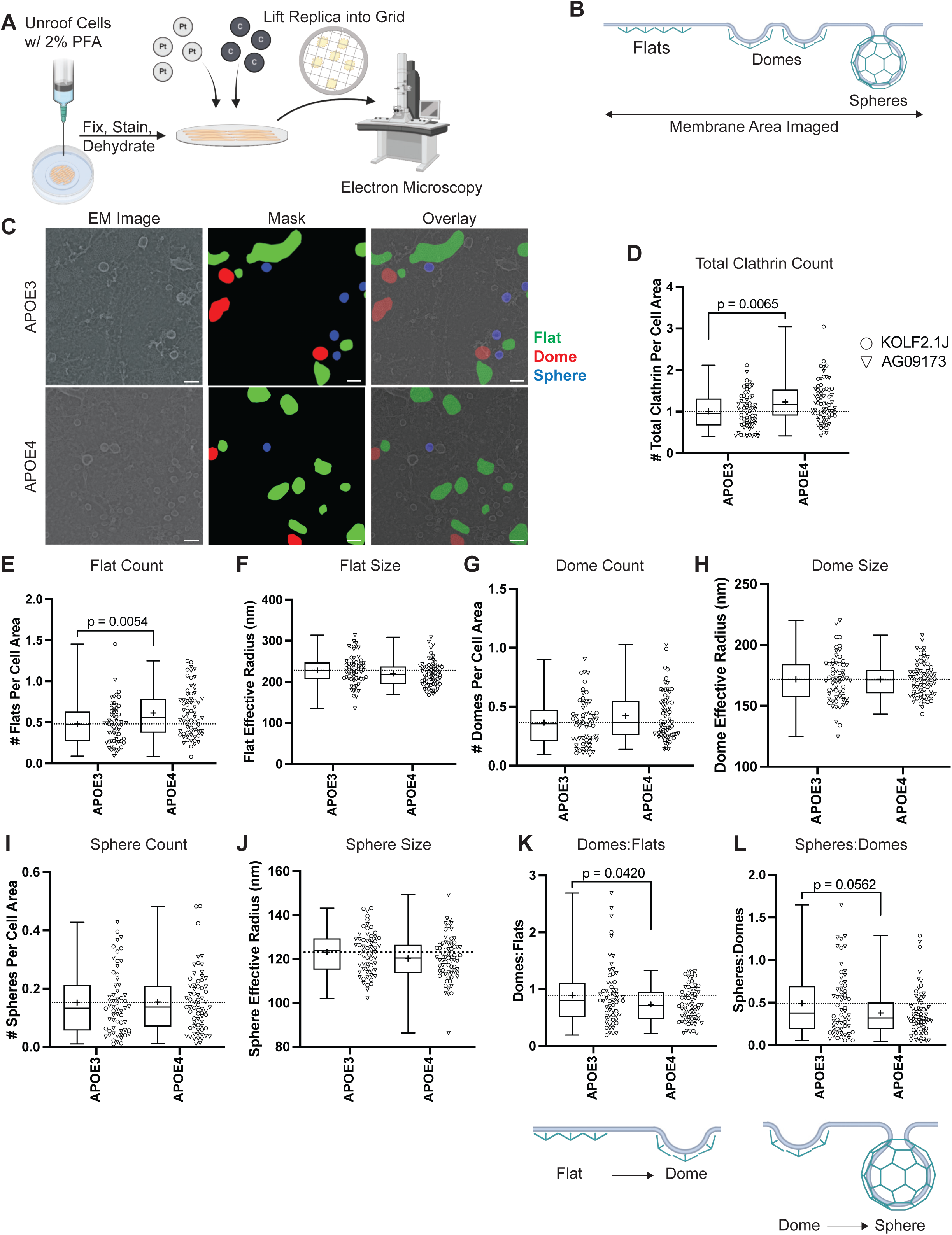
*APOE4* astrocytes display stalled clathrin-coated pit maturation. **A)** Schematic of unroofing and platinum replica electron microscopy (PREM) workflow. **B)** Diagram depicting a model of clathrin-mediated membrane curvature. **C)** PREM example images from *APOE3* and *APOE4* astrocytes containing segmented masks for flat, domed, and spherical clathrin structures. Scale bar is 100 nm. Quantification of **D)** number total clathrin structures, **E)** number of flat structures, **F)** average area of flat structures, **G)** number of dome structures, **H)** average dome radius, **I)** number of spherical structures, and **J)** average sphere radius. Quantification of **K)** Ratio of Domes:Flats and **L)** Spheres:Domes (structure count) per cell. N=61-68 cells per genotype, across two iPSC lines (KOLF2.1 O, AG09173 ∇). Diagrams for quantification are depicted with both box and whisker plots, minimum to maximum with line at median, cross (+) indicates the mean of each data set. Individual cell data is shown by column scatter plot with parental iPSC line indicated. Unpaired t-test used for statistical analysis.

Our data suggest that the *APOE4* astrocytes have clathrin lattice maturation defects at the structural level. Despite the increase in total clathrin structures at the plasma membrane of *APOE4* astrocytes, there is decreased ability to mature clathrin flats into downstream domed and spherical morphologies that drive the budding of clathrin-coated endocytic vesicles into the cytoplasm. This finding suggests that *APOE* genotype impacts clathrin lattice maturation and membrane curvature generation in astrocytes.

### *APOE4* decreases astrocytic endocytosis capacity in serum free culture conditions

We next asked whether the *APOE4-*associated defect in CCP maturation translates into impaired endocytic function. *APOE4-*associated endocytic dysfunction has been previously reported in astrocytes cultured in serum-containing media^7,12^.To explore whether the perturbed clathrin lattice maturation dynamics we observed translate into endocytic dysfunction under serum-free conditions, we used flow cytometry and immunocytochemistry to measure the uptake of a fluorescently conjugated transferrin ligand alongside characterizing levels of EEA1 (early endosomal antigen 1)-positive endosomes (Figure 2A). We observed a decreased uptake of the fluorescently labeled transferrin ligand (Figure 2B, 2D, and S2A, S2B) as well as decreased levels of EEA1 (early endosomal antigen 1)-positive early endosomes (Figure 2C, 2E and S2C, S2D) in *APOE4* astrocytes relative to their *APOE3* counterparts. This suggests that endocytic defects in *APOE4* astrocytes are linked with impaired CCP maturation and these defects are present in serum-free culture conditions.

**Figure 2:**
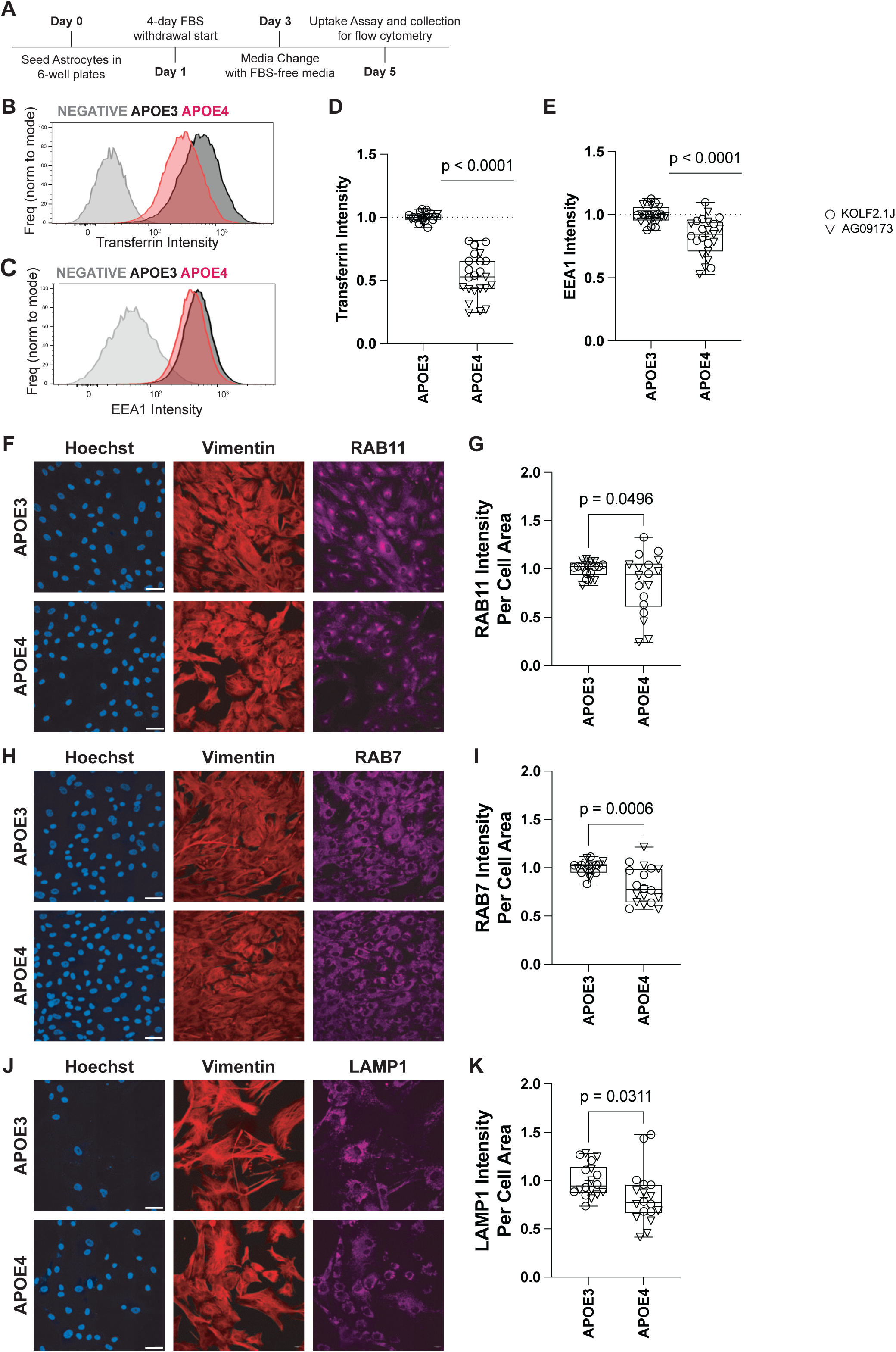
*APOE4* astrocytes exhibit endocytic defects compared to *APOE3* astrocytes in serum free culture conditions. **A)** Schematic for iPSC-derived astrocyte culture in serum-free media before endocytic characterization via flow cytometry. Representative histograms from flow cytometry quantification of **B)** fluorescently conjugated transferrin uptake and **C)** EEA1 intensity in *APOE4* and *APOE3* astrocytes from two different iPSC-lines. **D)** Box and whisker plot (minimum to maximum values with line at median) for quantification of transferrin uptake and **E)** EEA1 intensity, N=23 flow cytometry runs per genotype (11 for KOLF2.1J, 12 for AG09173), Wilcoxon signed-rank test. Representative images for **F)** RAB11 positive recycling endosomes with **G)** quantification, **H)** RAB7 positive late endosomes with **I)** quantification, and **J)** LAMP1 positive lysosomes with **K)** quantification. N=18 wells per genotype, across two iPSC lines (KOLF2.1 O, AG09173 ∇). Box and whisker, minimum to maximum with line at median, cross (+) denotes mean. Unpaired t-test used for statistical analysis. All scale bars are 50 µm.

Next, to determine whether defects in the early endocytic pathway propagated to later stages of endocytosis, we compared immunocytochemistry staining of RAB11-positive recycling endosomes (Figure 2F, 2G), RAB7-positive late endosomes (Figure 2H, 2I), and LAMP1-positive lysosomes (Figure 2J, 2K) between the *APOE3* and *APOE4* astrocytes. Compared to *APOE3* astrocytes, *APOE4* astrocytes showed decreased levels of all three downstream endosomal compartment markers (RAB11, RAB7, LAMP1) indicating dysfunction throughout the endocytic pathway. This contrasts previously reported findings from experiments performed in serum-containing media^12^ highlighting the importance of culture conditions when modeling human biology. Together, these data show that *APOE4-*associated defects in CCP maturation are linked to disrupted clathrin-mediated endocytosis and impair capacity at all stages of the endosome-lysosome pathway.

### *APOE4* alters plasma membrane lipid composition in astrocytes

Our findings indicate that *APOE4* disrupts clathrin-coated pit maturation at the plasma membrane, leading to impaired endocytosis. Given that CCP generation is highly sensitive to the mechanical properties of the plasma membrane ^29^ ^30^, we next investigated whether expression of *APOE4* alters membrane lipid composition and tension in astrocytes.

We examined the levels and molecular composition of plasma membrane lipids (phosphatidylserine, PS; phosphatidylinositol, PI; phosphatidylglycerol, PG; phosphatidylethanolamine, PE; phosphatidylcholine, PC; phosphatidic acid, PA; and sphingomyelin, SM) from publicly available whole cell lipidomics datasets on iPSC-derived *APOE3* and *APOE4* astrocytes^31^. Although the overall abundance of most phospholipid classes was comparable between *APOE3* and *APOE4* astrocytes, *APOE3* astrocytes were enriched in phosphatidylserine (PS) and sphingomyelin (SM), lipids that impact membrane curvature^29,30,32^. When we analyzed these data to characterize chain length and saturation of the fatty acids of membrane lipid species, we observed that across all membrane-enriched lipid classes, the fatty acid chain lengths were shorter and more saturated in *APOE4* astrocytes than *APOE3* astrocytes (Figure 3A, 3C). This shift towards shorter, saturated fatty acids in *APOE4* astrocytes was even more pronounced when examining the most abundant membrane lipid, phosphatidylcholine (PC; Figure 3B, 3D). These observations are consistent with reports from human brain lipidomics which measure lower levels of unsaturated plasma membrane lipids in disease brains relative to control brains^33^.

**Figure 3:**
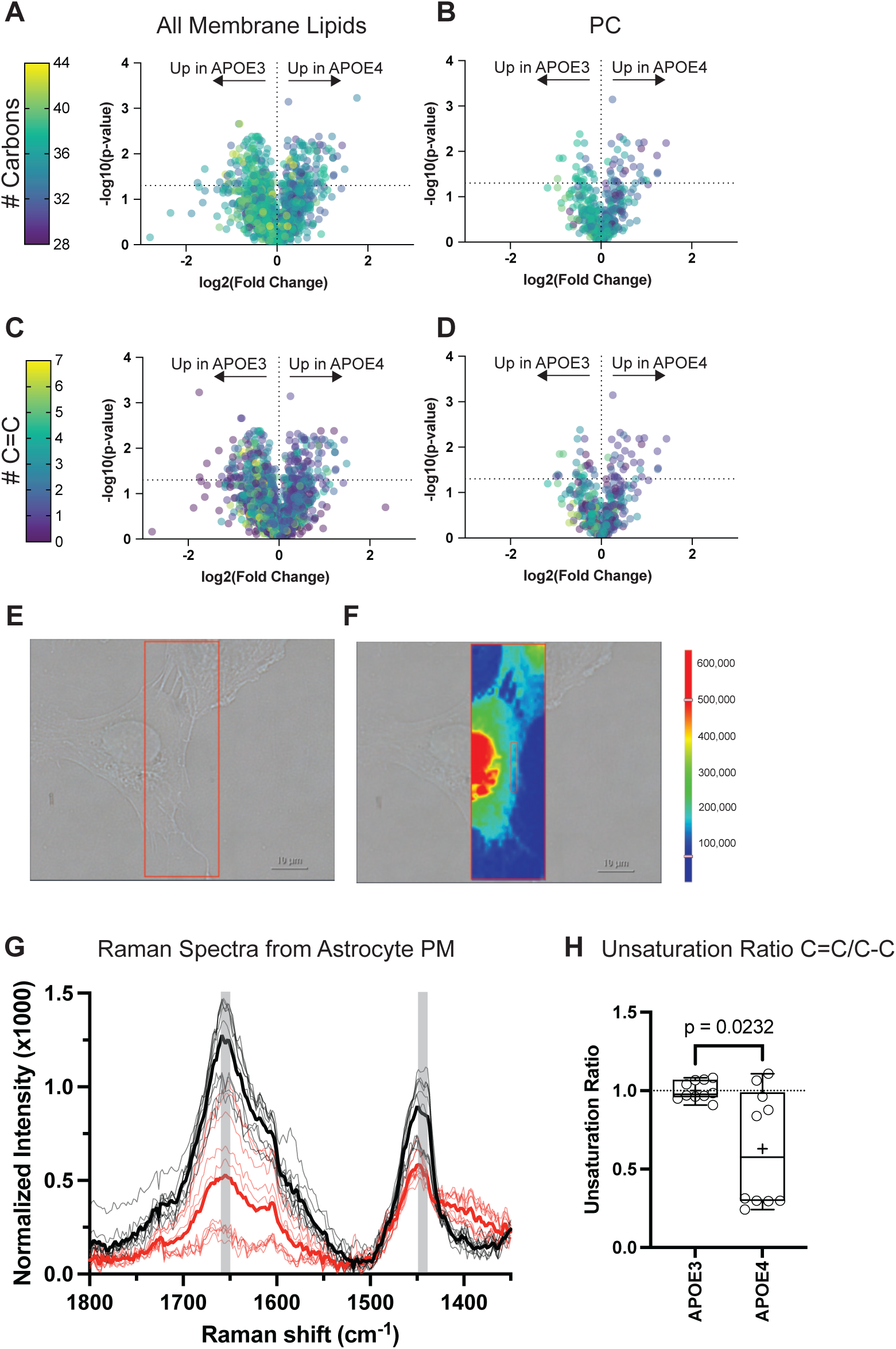
*APOE4* alters membrane lipid species composition in human iPSC-derived astrocytes. **A)** Volcano plot representing the length of the fatty acid chains from seven (PS, PI, PG, PE, PC, PA, and SM) membrane-enriched lipid species. **B)** Volcano plot representing the length of the fatty acid chains from PC only. **C)** Volcano plot representing the number of carbon double bonds (unsaturation) in fatty acid chains from the combined seven common membrane lipid species. **D)** Volcano plot representing the number of carbon double bonds (unsaturation) in fatty acid chains from PC only. Volcano plot dots represent an individual lipid member of the denoted class from 3 different iPSC lines combined (BIONi-037A, KOLF2.1J #1, and KOLF2.1J #2). **E)** Example brightfield image of an iPSC-derived astrocyte captured for Raman spectroscopy analysis, along with **F)** a representative Raman spectroscopy-based chemigram, scale shows raw intensity values, white box highlights an example of a region of interest for membrane analysis. **G**) Raman spectrum from 1350 cm^-1^ to 1800 cm^-1^ (region of interest for C=C double bonds and C-C single bonds) from N=10 membrane regions from KOLF2.1J *APOE3* astrocytes (grey, solid black = *APOE3* average) and from N=10 membrane regions from KOLF2.1J *APOE4* astrocytes (pink, solid red = *APOE4* average). Shaded areas represent peak range associated with C=C double bonds (1652-1660 cm^-1^) and C-C single bonds (1440-1448 cm^-1^). **H)** Unsaturation ratio for *APOE3* and *APOE4* plasma membrane regions determined from Raman spectra. Box and whisker, minimum to maximum with line at median, cross (+) indicates mean. Unpaired t-test used for statistical analysis.

We next used Raman spectroscopy to directly measure membrane lipid saturation in our *APOE3* and *APOE4* astrocytes (Figure 3E, 3F, S3A). Membrane regions from *APOE3* astrocytes generated spectra with stronger signals for C=C double bonds than *APOE4* spectra (Figure 3G). To quantify this, we determined the ratio between the area under the curve in the C=C double bond region and the C-C single bond region to generate an unsaturation ratio (Figure 3H). This measurement further confirmed that *APOE4* astrocytic plasma membranes have more saturated (lower ratio) lipid species compared to their *APOE3* counterparts.

### *APOE4* astrocytes have increased membrane tension

Since fatty acid saturation influences the biophysical properties of the plasma membrane^29,30,32–34^, we measured membrane tension directly in *APOE3* and *APOE4* astrocytes using the Flipper-TR fluorescent cell membrane tension probe (Figure 4A). We then quantified the fluorescence lifetime decay via fluorescence lifetime imaging microscopy (FLIM)^35^ (Figure 4B). *APOE4* astrocyte plasma membranes had longer fluorescence lifetimes compared to *APOE3* astrocytes, indicating increased membrane tension in *APOE4* astrocytes (Figure 4B,C). Together, these results demonstrate that *APOE4*-associated increased fatty acid saturation in the plasma membrane translate to altered membrane tension, providing a biophysical basis for impaired generation of clathrin curvature.

**Figure 4:**
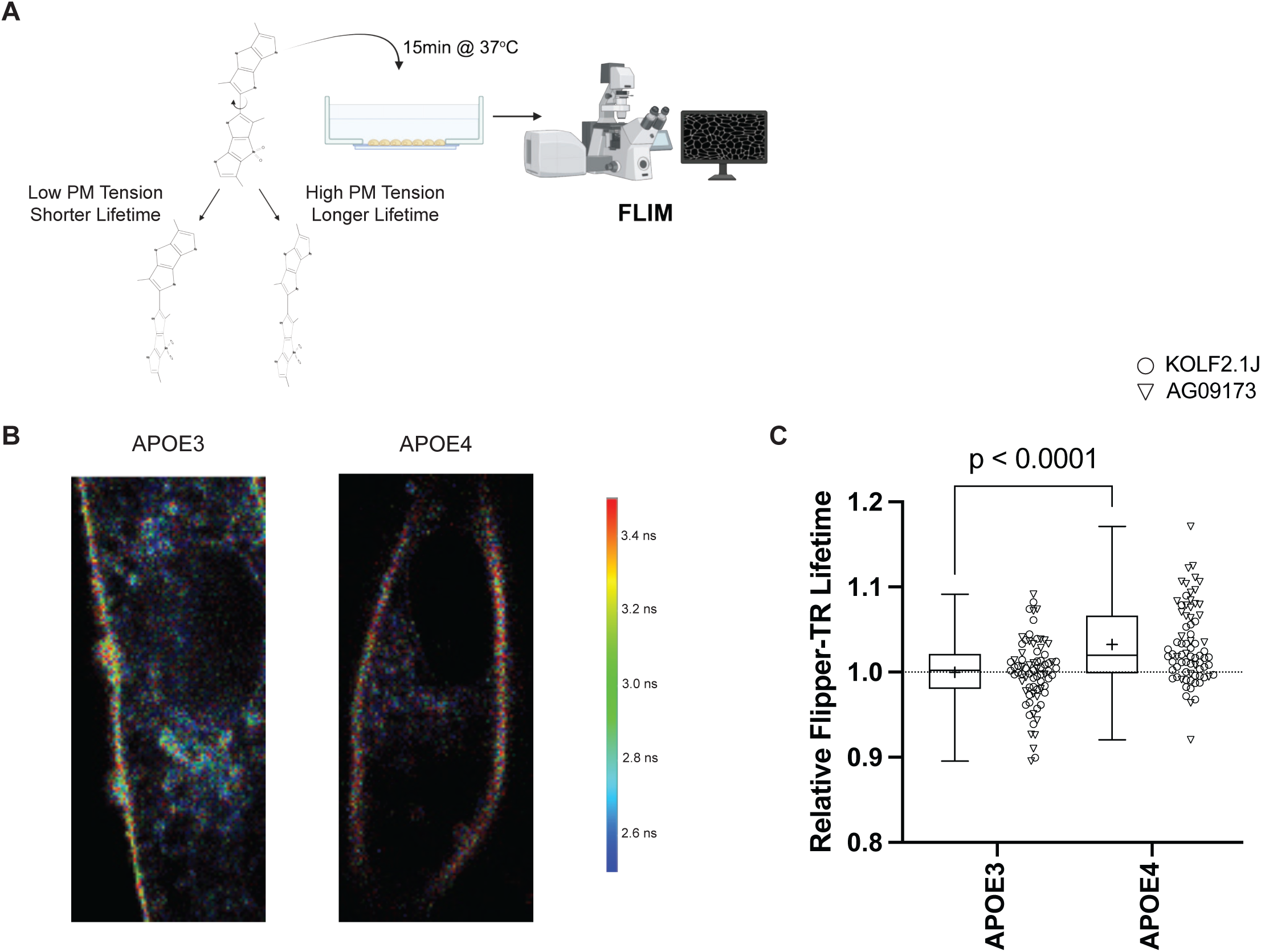
*APOE* genotype alters plasma membrane tension in astrocytes. **A)** Schematic of Flipper-TR^®^ plasma membrane tension quantification assay in astrocytes, including diagram of low plasma membrane tension conformation and high plasma membrane tension conformation changes in Flipper-TR^®^ structure. **B)** A representative set of FastFLIM heatmap images of *APOE3* and *APOE4* astrocytes. **C)** Phasor analysis results for Flipper-TR^®^ lifetime decay in *APOE3* and *APOE4* astrocytes (normalized to *APOE3*). N=76-78 images per genotype (KOLF2.1 O, AG09173 ∇), box and whisker, minimum to maximum with line at median, cross (+) denotes mean. Unpaired t-test used for statistical analysis.

### Increasing *INPP5D* expression rescues early endocytosis defects in *APOE4* astrocytes

Next, we asked whether increasing the expression of proteins associated with endocytosis could overcome the early endocytic defect in *APOE4* astrocytes. Previous work demonstrated that increased expression of the AD risk gene *PICALM*, a clathrin adaptor protein, restores endocytosis in *APOE4* astrocytes^12^. Given that multiple endocytic proteins share membrane and clathrin binding functionalities, we hypothesized that providing exogenous levels of individual endocytic proteins might similarly rescue *APOE4*-associated defects^36–40^. To test this, we systematically increased expression of eight endocytic proteins, *SNAP91*, *INPP5D*, *BIN1*, and five individual AP2 complex subunits, in *APOE4* astrocytes via lentiviral transduction (Figure S4A-C). This set of proteins included both interactors of PICALM as well as AD risk factors that are also involved in endocytosis^3^. Endocytosis in cells transduced with each of these candidate proteins was compared to their respective effects in *APOE4* astrocytes transduced with a lentivirus expressing a control protein, a non-fluorescent eGFP mutant (Y66L, “darkGFP”). Exogenous protein expression was quantified by the detection of a C-terminal HA tag (Figure S4C) and was found to be expressed at comparable or lower levels than the darkGFP control for all proteins examined (Figure S4D, S4E). Following lentiviral transduction, we assessed EEA1 levels and uptake of fluorescently tagged transferrin using flow cytometry (Figure 5A, 5B).

**Figure 5:**
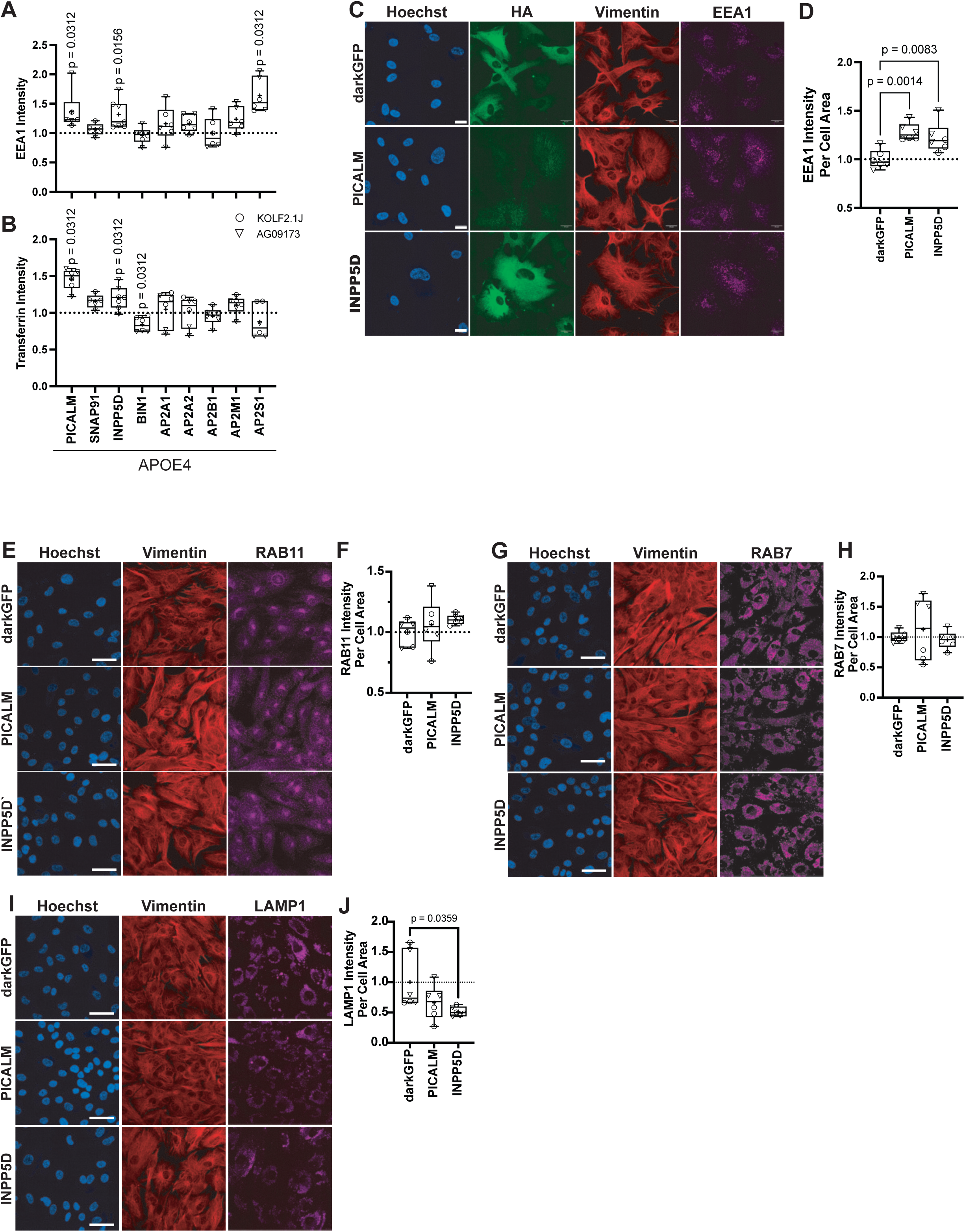
Increasing *INPP5D* or *PICALM* expression rescues *APOE4*-mediated endosomal dysfunction in astrocytes. Box and whisker plot (minimum to maximum values with line at median) for quantification of multiple flow cytometry runs for **A)** EEA1 and **B)** internalized transferrin intensity, N=5-7 independent flow cytometry runs per endocytosis protein, across two iPSC lines (KOLF2.1 O, AG09173 ∇), flow cytometry means normalized to *APOE4* astrocytes transduced with control *darkGFP* expressing virus from same run. Wilcoxon signed-rank test. **C)** Representative immunocytochemistry images from *APOE4* astrocytes with increased *darkGFP*, *PICALM*, and *INPP5D* expression (scale bar 20 µm) with **D)** EEA1 intensity quantification. N=6 independent experiments per condition per iPSC line (KOLF2.1 O, AG09173 ∇). One-way ANOVA with multiple comparisons used for statistical analysis. Representative immunocytochemistry images from *darkGFP*, *PICALM*, and *INPP5D* overexpressing *APOE4* astrocytes for **E)** RAB11 positive recycling endosomes with **F)** quantification, **G)** RAB7 positive late endosomes with **H)** quantification, and **I)** LAMP1 positive lysosomes with **J)** quantification. Scale bars are 50 µm.

In addition to *PICALM*, increasing expression of *INPP5D* (SHIP1) and *AP2S1* (AP2 s subunit) increased EEA1 levels in *APOE4* astrocytes (Figure 5A). Only *PICALM* and *INPP5D*, not *AP2S1* overexpression increased the uptake of fluorescently tagged transferrin in *APOE4* astrocytes while *BIN1* overexpression impaired transferrin uptake (Figure 5B). Immunocytochemistry confirmed that increased expression of either *PICALM* or *INPP5D* can alleviate endocytic dysfunction in *APOE4* astrocytes (Figure 5C, 5D). We also used immunocytochemistry to assess whether perturbations to downstream trafficking pathways were also rescued by increasing *PICALM* or *INPP5D* expression. Increasing either *PICALM* or *INPP5D* expression had no detectable effect on either RAB11-positive recycling endosomes (Figure 5E, 5F) or RAB7-positive late endosomes (Figure 5G, 5H). Increased *INPP5D* expression did result in reduced LAMP1-positive lysosome staining compared to darkGFP (Figure 5I, 5J). These findings identify *INPP5D* as a second Alzheimer’s disease risk gene that can restore early endocytosis defects in *APOE4* astrocytes.

### *INPP5D* restores clathrin-coated pit maturation in *APOE4* astrocytes

Given that increasing levels of *INPP5D* or *PICALM* functionally restore early endocytic defects in *APOE4* astrocytes, we further characterized the effects of increased *INPP5D* or *PICALM* expression on membrane tension or clathrin morphology. We used fluorescence-activated cell sorting to isolate pure populations of *APOE4* astrocytes that have been stably transduced by lentivirus to express *darkGFP*, *PICALM*, or *INPP5D*. Then, we measured the lifetime of the membrane tension sensor, Flipper-TR, in these transduced cells. We observed that only exogenous expression of *PICALM*, not *INPP5D*, decreased membrane tension in *APOE4* astrocytes (Figure 6A). This suggests that the two genetic modifiers of endocytosis, *PICALM* and *INPP5D*, may operate through distinct mechanisms.

**Figure 6:**
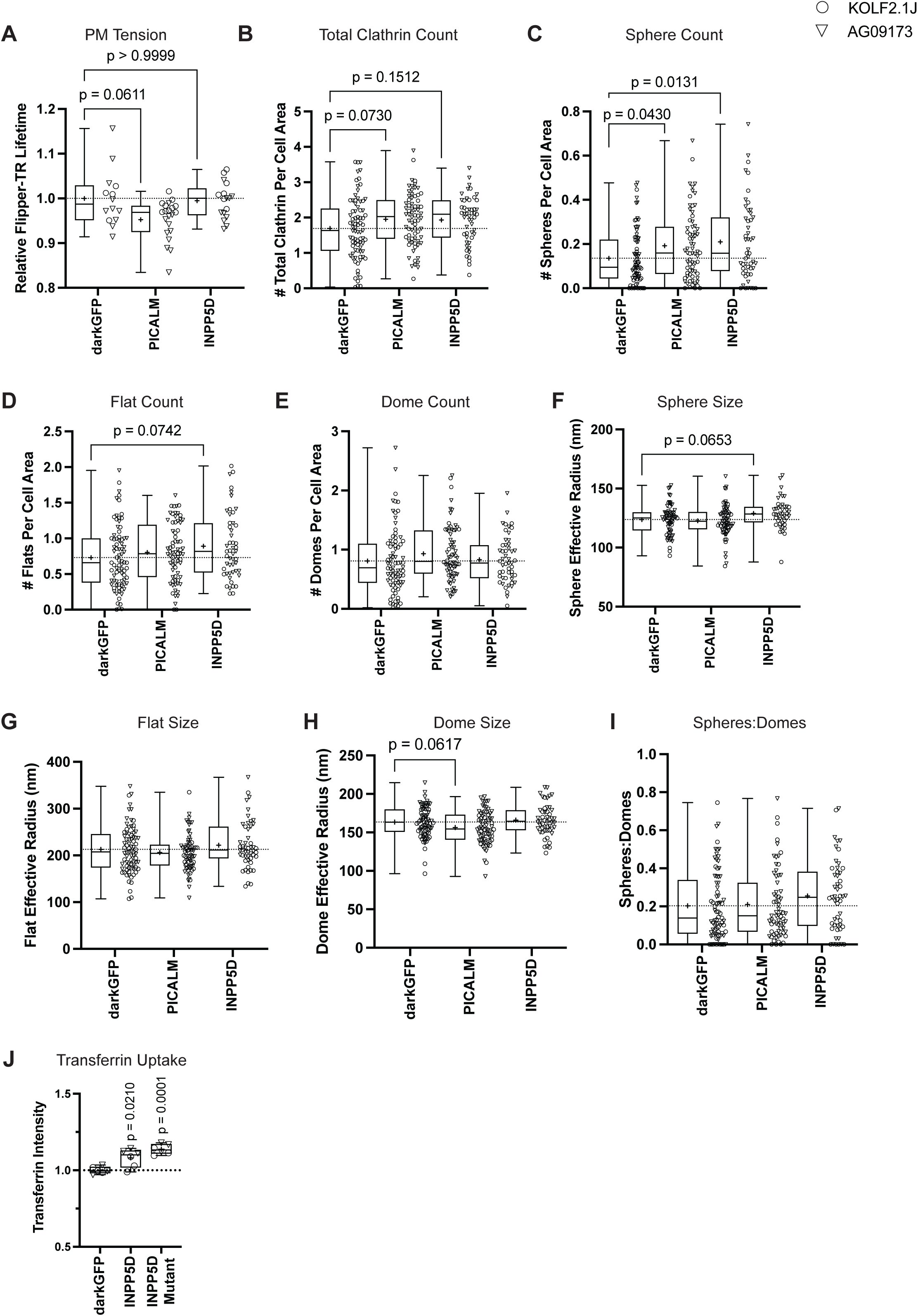
Increased *PICALM* and *INPP5D* expression alter plasma membrane tension and clathrin morphology in *APOE4* astrocytes. **A)** Phasor analysis results for Flipper-TR lifetime in *APOE4* astrocytes expressing *darkGFP*, PICALM, or INPP5D (normalized to darkGFP). N=14-21 images per adaptor protein, across two iPSC isogenic lines (KOLF2.1 O, AG09173 ∇). Box and whisker plot (minimum to maximum values with line at median), cross (+) denotes mean, One-way ANOVA with multiple comparisons used for statistical analysis. PREM quantification of number of **B)** total clathrin, **C)** spherical, **D)** flat, and **E)** dome structures per cell area. **F)** Average sphere radius, **G)** flat radius, and **H)** dome radius per cell. **I)** Ratio of Spheres:Domes (structure counts) per cell and N=44-81 cells per condition, across two iPSC isogenic lines (KOLF2.1 O, AG09173 ∇). Box and whisker, minimum to maximum with line at median, cross (+) denotes mean. One-way ANOVA with multiple comparisons used for statistical analysis. Box and whisker plot (minimum to maximum values) for quantification of multiple flow cytometry runs for **J)** internalized transferrin intensity, N=6 independent flow cytometry runs per condition, across two iPSC lines (KOLF2.1J, AG09173), flow cytometry means normalized to *APOE4* astrocytes transduced with control *darkGFP* values from same run, Wilcoxon signed-rank test.

We then examined clathrin architecture using PREM. Increasing expression of both *PICALM* and *INPP5D* showed a modest, though statistically insignificant, trend towards a higher number of total clathrin structures at the plasma membrane relative to control (Figure 6B, S5A). When we classified these structures as flats, domes, and spheres, we noticed that the number of spherical clathrin structures was significantly higher in cells with either increased *INPP5D* or *PICALM* expression (Figure 6C); the number of flat clathrin structures trended higher in cells with increased *INPP5D* expression (Figure 6D) while the number of domes was not significantly different across the conditions (Figure 6E). Only increased *INPP5D*, not *PICALM* expression, showed a trend towards increased size of spherical clathrin structures (Figure 6F) and no significant changes were observed in flat or dome clathrin sizes with *INPP5D* (Figure 6G, 6H). Interestingly, increased *PICALM* expression led to a non-significant decrease in flat and dome sizes (Figure 6G, 6H). Increasing *INPP5D* expression did not influence dome:flat ratio (Figure S5B) but did show a trend towards increased sphere:dome ratio suggesting a slight increase in efficiency of conversion between domes and spheres (Figure 6I).

These data on membrane tension and clathrin structure support distinct rescue mechanisms. *PICALM* decreases membrane tension and increases clathrin structures at the membrane whereas *INPP5D* does not modify membrane tension but also increases clathrin structures and increases the domed to spherical clathrin structure conversion.

*INPP5D* encodes a lipid phosphatase that converts PI(3,4,5)P₃ to PI(3,4)P₂, a reaction important for endocytosis^41^. To determine whether its catalytic activity is required for rescue, we overexpressed a phosphatase-dead INPP5D mutant (P668A/D672A/R673G) in *APOE4* astrocytes^42^. Like wild-type INPP5D, the mutant increased transferrin uptake (Figure 6J). Since mutant expression levels were lower than wild-type *INPP5D* (Figure S5C, S5D), these results suggest that INPP5D-mediated rescue of *APOE4*-impaired endocytosis could be independent of its phosphatase activity.

### Increasing *PICALM* and *INPP5D* expression alters disease-relevant cellular phenotypes

Lipid remodeling and altered membrane mechanics leading to endosomal dysfunction represent early cellular consequences of *APOE4* on AD. Given our identification of potent modifiers of early endocytic capacity, we next asked whether modulation of this same pathway impacts other *APOE4-*associated AD-relevant astrocyte phenotypes. Lipid droplet (LD) accumulation and dysregulated lipid homeostasis is a well-established feature of *APOE4* astrocytes and AD pathology^31,43–47^. Recent work has shown that AD associated variants that decrease *PICALM* and *INPP5D* expression increase the accumulation of lipid droplets in microglia^48,49^. We explored whether increasing expression of *PICALM* and *INPP5D* changed lipid droplet accumulation in astrocytes. We discovered that increasing expression of *INPP5D* in *APOE4* astrocytes reduced total amount and size of LDs (Figure 7A-C). These findings suggest that *INPP5D* and its genetic interactions with *APOE* genotypes may have broad effects on AD disease-associated phenotypes beyond endocytosis.

**Figure 7:**
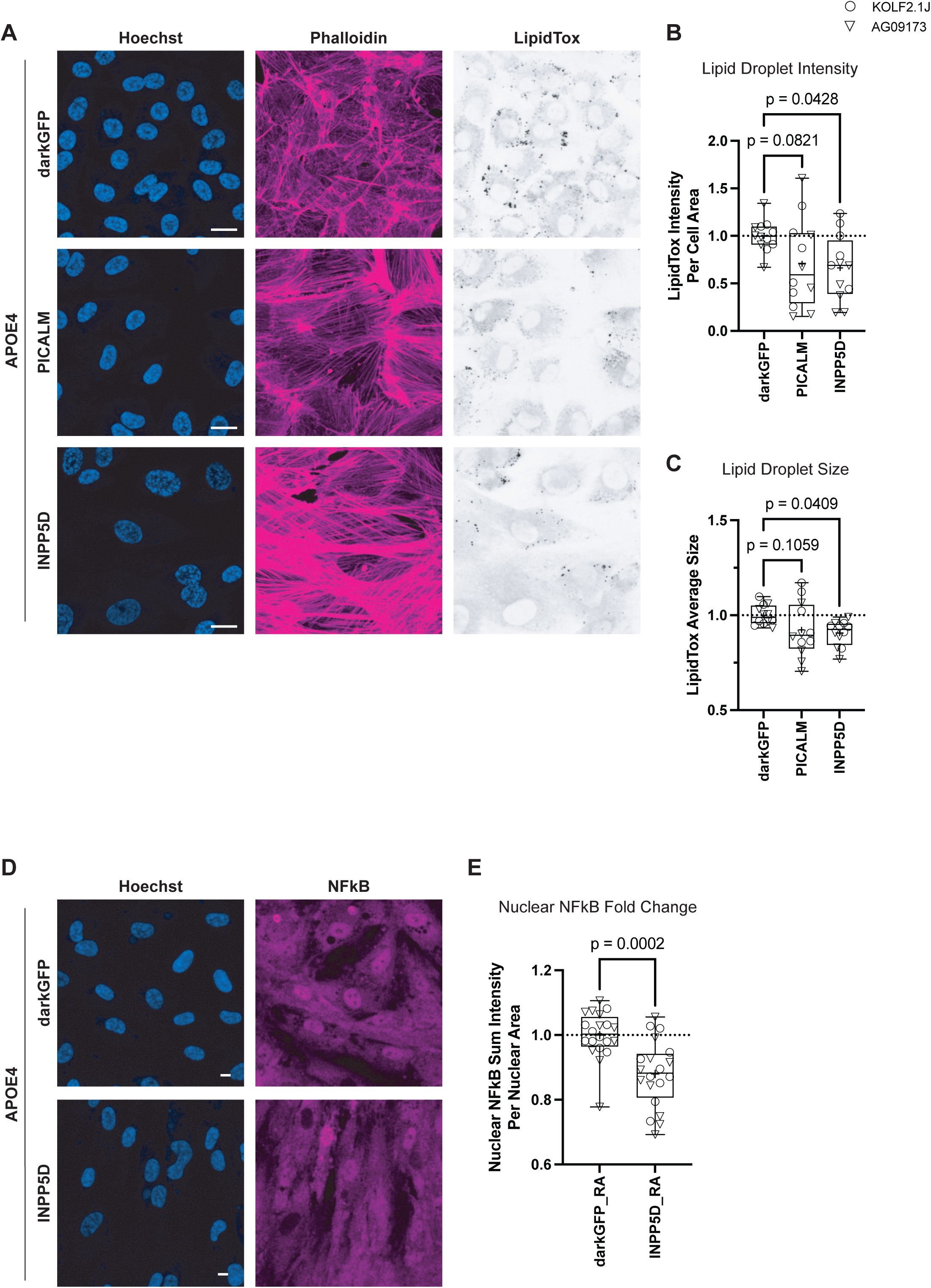
*INPP5D* (SHIP1) decreases lipid droplet burden and immune activity of *APOE4* astrocytes. Representative images for **A)** LipidTox staining, with quantification for **B)** lipid droplet intensity and **C)** lipid droplet size, N=12 wells per condition, across 2 iPSC isogenic lines (KOLF2.1 O, AG09173 ∇). Box and whisker, minimum to maximum with line at median, cross (+) denotes mean. Scale bar is 20 µm) One-way ANOVA with multiple comparisons used for statistical analysis. Representative images for **D)** reactive astrocyte NFκB localization (Scale bar is 50 µm), with quantification for **E)** relative NFκB nuclear signal fold change (normalized to vehicle treatment), N=20 independent activation replicates per condition, across 2 iPSC isogenic lines (KOLF2.1 O, AG09173 ∇). Box and whisker, minimum to maximum with line at median, cross (+) denotes mean. One-way ANOVA with multiple comparisons used for statistical analysis.

Neuroinflammation and specifically activation of astrocytes is also a key feature of AD pathology^50,51^. Moreover, perturbations to lipid metabolic pathways have been linked to activation in various glial cell types^23,31,52,53^. We therefore assessed the translocation of the inflammation-associated transcription factor, NF-κB, to the nucleus following astrocyte activation by delivering TNF-α, IL-1α, and C1q in the cell culture media^54^. Exogenously increasing *INPP5D* expression reduced activation-associated NF-κB nuclear localization in *APOE4* astrocytes (Figure 7D,E). This finding suggests a broadly protective role for *INPP5D* expression in *APOE4* astrocytes beyond its role in endocytosis.

Together, these findings show that modulating endocytic proteins in *APOE4* astrocytes has effects beyond restoration of endocytic trafficking and influences metabolic and inflammatory phenotypes relevant to Alzheimer’s disease.

## Discussion

Genome-wide association studies and functional work have implicated endocytic pathway genes and dysfunction in late-onset Alzheimer’s disease, yet the mechanistic underpinnings of these findings are poorly understood^4,6,7,12^. Here, using isogenic human iPSC-derived astrocytes, PREM, endocytic uptake assays, and plasma membrane biophysics, we discovered that *APOE4* astrocytes have altered membrane lipid composition that possibly explains the observed increase in *APOE4* astrocyte plasma membrane tension and links to defects in clathrin curvature and endocytosis. Our findings align well with lipidomic findings from dementia patient brains exhibiting decreased plasma membrane polyunsaturated fatty acids (PUFAs) compared to age-matched, healthy controls^33^.

Since *APOE* encodes a lipid transport protein, our findings suggest that plasma membrane lipid remodeling may represent a proximal consequence of the *APOE4* genotype. By linking *APOE4*-dependent changes in fatty acid composition to increased membrane tension and impaired clathrin curvature, we propose that altered membrane mechanics may represent one of the many ways in which *APOE4-*associated perturbations to lipids impact cell biology. This finding contributes to a greater hypothesis emerging from the field, that lipid disruptions, due to their pleiotropic consequences, may significantly contribute to neurodegenerative disease risk^14,23,31,33,46,52,53,55,56^.

We identified *INPP5D* as an AD risk gene capable of restoring endocytosis in *APOE4* astrocytes and show that it acts via a mechanism distinct from a previously identified endocytic modulator, *PICALM*^12^. Increasing *PICALM* expression lowers membrane tension and increases clathrin availability at the membrane while *INPP5D* promotes clathrin curvature progression without altering tension. Our experiments showed that INPP5D’s effect is not dependent on its phosphatase activity, suggesting that its rescue may arise from other functions, perhaps its role as an endocytic scaffold^57^. AD risk-associated variants in both *INPP5D* and *PICALM* decrease expression of the respective proteins while AD protective variants have been shown to increase protein expression^48,49,58–62^. This finding aligns with our discovery of increasing *INPP5D* and *PICALM* expression reducing *APOE4-*associated defects. In contrast, increased expression of another AD risk gene, *BIN1*, did not rescue endocytosis in *APOE4* astrocytes. This suggests a possible divergence in disease risk and protective mechanisms between different AD-associated genetic variants.

Since membrane remodeling and endocytic dysfunction can impact a broad range of cellular functions, we tested whether correcting these defects impacts disease-associated astrocyte phenotypes. Increasing *INPP5D* expression reduced lipid droplet accumulation and the reactivity of astrocytes in response to cytokine treatment. This finding suggests that modulating membrane dynamics and endosomal trafficking can influence downstream metabolic and inflammatory phenotypes linked to *APOE4*, though future work will help untangle the molecular cascades responsible for this link. Together, these results advance a model in which multiple AD risk genes modulate membrane tension, regulate clathrin dynamics, and impact disease phenotypes in astrocytes.

Astrocytes are integrative cell types within the central nervous system that maintain homeostasis through cell autonomous and non-autonomous processes^51^. Impaired clathrin-mediated uptake may impact pathways central to astrocyte function including clearance of extracellular material, neuron-glial and glial-glial communication, receptor turnover, and cytokine signaling. Therefore, changes to astrocytic endocytosis, though not disease causing on their own, could influence disease susceptibility through impaired cell health. Disrupted membrane curvature and endocytosis may alter synaptic homeostasis and neuronal excitability in a cell non-autonomous manner. In another previous study, an astrocyte-specific knockout of saturated lipid elongase *ELOVL1* rendered activated astrocytes less toxic to their neuronal neighbors, suggesting that modulation of lipid pathways can impact cell nonautonomous biology^63^. Future work will investigate the cell non-autonomous consequences of altering astrocytic endocytosis.

Although this study focuses on the plasma membrane, organelle membranes are defined by distinct lipid compositions that govern their function. It remains possible that *APOE4*-associated lipid changes extend to endosomal, ER, or mitochondrial membranes, contributing to previously reported defects in these compartments^64–66^. Defining whether altered cellular membrane mechanics represents a global feature of *APOE4* dysfunction will require further investigation. In summary, our findings link altered membrane mechanics and clathrin curvature defects in *APOE4* astrocytes and reveal functional convergence with additional AD risk genes at the level of endocytosis. By linking genetic risk to clathrin structures, this work provides a mechanistic framework for understanding how AD-associated genes may influence cellular vulnerability and disease.

## Limitations

While we identify altered membrane mechanics in *APOE4* astrocytes, we do not explore whether similar biophysical changes occur across additional CNS cell types or *in vivo*. Our data support the functional convergence of *APOE4*, *INPP5D*, and *PICALM* on endocytosis, but we do not yet know whether these same endocytic disruptions are a part of the risk biology of other AD-associated variants. Our rescue experiments relied on exogenously increasing expression using lentivirus. The efficacy of endogenous modulation of these pathways remains to be determined. In this study we explore *APOE3* and *APOE4* biology, but other *APOE* variants exist. Future work will explore the effects of *APOE2* and other rare *APOE* variants like *APOE-R136S* on plasma membrane dynamics.

## Materials and Methods

All lentiviruses and antibodies used are detailed in Supplemental Table 1.

### Astrocytes differentiation and maintenance

Two isogenic *APOE* iPSC sets were used for all experiments. These lines were generated from parental lines AG09173 (Coriell) and KOLF2.1J (The Jackson Laboratory; JAX) and acquired via MTA from MIT or purchased directly from JAX. These iPSCs were karyotyped prior to differentiation and were differentiated into astrocytes following a previously published small-molecule guided protocol^7^. Briefly, human induced pluripotent stem cells were derived into neural progenitor cells using the StemDiff SMADi Neural Induction kit (StemCell Technologies, 08582) as per manufacturer’s instructions (monolayer protocol). Then neural progenitors were differentiated into astrocytes using supplementation of BMP4 and FGF2 for 4 weeks. These cells were then sorted for an ACSA-1 positive population (Miltenyi Biotech 130-123-555), grown for two weeks in Astrocyte Medium (ScienCell 1801), and re-sorted for a purely ACSA-1 positive population. Pure iPSC-derived astrocytes were cultured using Astrocyte Medium (AM) containing 2% fetal bovine serum (FBS) (ScienCell 1801) on tissue culture treated 6- and 12-well plates, and 10-cm and 15-cm dishes. iPSC-derived astrocytes were passaged using TrypLE^TM^ (ThermoFisher Scientific, 12604039) once they reached confluency and split to a density of 1x10^6^ cells per 10 cm dish. All iPSC-derived astrocytes were used for experiments prior to passage 11.

### Platinum replicate electron microscopy

Human-derived iPSC astrocytes were seeded directly onto round #1.5 glass coverslips 25mm (Warner Instruments 64-0715). Cells were grown on coverslips in FBS-free AM with astrocyte growth supplement (AGS) and penicillin for 18-20 hours. (ScienCell 1801). At this point, cells were removed from the incubator for unroofing. For unroofing, cells were rinsed in stabilization buffer (30mM HEPES, 70mM KCl, 5mM MgCl2, 3mM EGTA at pH 7.4) for 10 seconds. Then cells were unroofed using our a lab-standardized, custom unroofing instrument, a machine with a 20-gauge needle spraying 2% PFA diluted in stabilization buffer at 0.7 bar of air pressure (Electron Microscopy Sciences 15710). After unroofing, cells were moved into fresh 2% PFA. Cells were covered and fixed at room temperature in the fresh PFA for 20 minutes. Next, cells were covered and further fixed at room temperature in 2% glutaraldehyde for 20 minutes (Electron Microscopy Sciences 16019). Cells were washed 3 times in water by moving cells to new wells with fresh water. Staining proceeded as follows: cells were placed in 0.1% tannic acid (Electron Microscopy Sciences, 1401-55-4) for 20 minutes, covered and at 4°C. Next, cells were washed 6 times in water by moving cells to new wells. Then cells were placed in 0.1% uranyl acetate for 20 minutes, covered and at 4C (Electron Microscopy Sciences 22400). Cells were washed at least 3 times in water. Next, dehydration proceeded as follows: cells were placed in increasing concentrations of EtOH for 4 minutes each (15%, 30%, 50%, 70%, 80%, 90%, 100% 3 times). Then cells were critically point dried using the autosamdri-815 (Tousimis Research Corporation) for 15 minutes. Coating proceeded as follows: coverslips were cut with a diamond knife and placed into the ACE-900 (Leica) where they were coated with 3nm of carbon and 5.5nm of platinum. Replicas were lifted from coverslips using hydrofluoric acid and then placed onto TEM grids (Ted Pella 01802-F). TEM imaging was performed with a JEM-1400 (JEOL) and montages generated via SerialEM software (University Colorado Boulder).

### Astrocyte lentiviral transduction

Confluent iPSC-derived astrocytes were detached using TrypLE^TM^ Express (ThermoFisher Scientific, 12604039) (10 minutes in a 37°C incubator) and pelleted by centrifugation (200xg for 3 min). 200,000 to 500,000 living cells were resuspended and plated using AM (ScienCell 1801) containing 2% FBS and 10 multiplicity of infection (MOI) of appropriate lentivirus. Lentiviral constructs were obtained from VectorBuilder, Inc. After 24 hours, transduced astrocytes were supplemented with fresh AM containing 2% FBS and further cultured for 48 hours. After a total of 72 hours of transduction, media was replaced with serum-free AM (without FBS).

### Endocytosis uptake assays

For microscopy, astrocytes were plated in square bottom 96-well imaging plates (ibidi, 89626-90) at 2,000-5,000 cells per well before start of 4 day serum (FBS) withdrawal and assaying post day 4. Astrocytes were incubated for 5 minutes at 37°C with fresh AM containing fluorescently conjugated transferrin (ThermoFisher, T35352) at 25 mg/mL. After incubation, astrocytes were directly placed onto ice and washed with cold DPBS (Quality Biological 114-057-101), before being fixed with 4% paraformaldehyde (Electron Microscopy Sciences EMT-5156). For flow cytometry, astrocytes were plated in 6 well, flat bottom dishes (Corning, 353046 at 150,000-200,000 cells per well before start of 4 day serum (FBS) withdrawal and assaying post day 4. Astrocytes were incubated for 5 minutes at 37°C with fresh AM containing fluorescently conjugated transferrin (ThermoFisher T35352) at 25 mg/mL. After incubation, astrocytes were directly placed onto ice and washed with cold DPBS (Quality Biological 114-057-101) and detached for collection using cold TrypLE^TM^ Express (ThermoFisher Scientific, 12604039) before fixing with 4% paraformaldehyde (Electron Microscopy Sciences EMT-5156) for 15 minutes at room temperature.

### Immunocytochemistry

5,000 to 7,500 astrocytes were plated per well of a square 96-well imaging plate (ibidi 89626-90), in AM containing 2% FBS (ScienCell 1801). After 24 hours, the cell culture medium was changed to serum-free AM and the cells were cultured in this medium for 4 days with media changes every other day. Cells were then fixed using 4% paraformaldehyde (Electron Microscopy Sciences, EMT-5156) in DPBS for 15 minutes at room temperature. Fixed cells were incubated in blocking buffer—1% normal donkey serum (NDS; Sigma, S30-100ML), 1% normal goat serum (NGS; Sigma, S26-100ML), 0.05% saponin (Sigma SAE0073), and 5% bovine serum albumin (BSA; Sigma, A7030-100G) in PBS at room temperature for 1 hour. Astrocytes were then incubated overnight at 4°C in blocking buffer containing primary antibodies. Primary antibody mixture was washed off with PBS at room temperature (3 times 5 minutes), before incubating with secondary antibodies for 1 hour at room temperature (in abovementioned blocking buffer composition). Cells were washed with PBS (3 times 5 minutes) before image acquisition with a Nikon CSU-W1 Spinning Disk Microscope at 20X or 40X magnification, using NIS Elements version 6.10.01 (Nikon).

### Raman spectroscopy of astrocyte plasma membrane regions

Samples used for Raman spectroscopy were transferred into μ-Dish 35 mm, high Glass Bottom microscopy dish (ibidi, 81158) and cultured (as described above), fixed with 4% paraformaldehyde in DPBS, and sealed with a round glass coverslip. Raman spectra were acquired using a DXR2xi Raman microscope (ThermoFisher Scientific) with 10 mW of 532 nm laser directed at the sample through a 100X oil confocal objective (N.A.=1.3 a working distance of 0.2 mm UPLanFLN Olympus America Inc.) at 2 s exposure time, through a 50 μm pinhole, for a pixel size between 200 nm and 1 μ, for the 100 to 3400 cm^-1^ spectra region. Chemical images were produced by the peak height selection of the assigned peaks using Thermo Fisher Scientific OMNICxi software (ThermoFisher Scientific), using a local selected baseline correction and displayed as a heatmap. Spectra from representative random regions of the cellular membrane were selected for spectral extraction. The edges of the spectra between 200 – 500 cm^-1^ and 3100 – 3400 cm^-1^ were eliminated, as they do not contain any relevant biological information. Next an automated background subtraction function was applied, followed by normalization to the sum and finally spectra were overlayed using Origin Pro 2021b (Origin Lab Corporation). An average spectrum was highlighted from at least five spectral regions of the membrane. The regions of interest were plotted and saved as Excel files and replotted in Graph Pad Prism (GraphPad Software, LLC) for further analysis and display. Unsaturation ratio was determined by dividing the peak are with a width of 1652 – 1660 cm^-1^ for the C=C double bond to 1440 – 1448 cm^-1^ for the C-C single bond^67,68^.

### Flow cytometry

Fixed iPSC-derived astrocytes were incubated in 100 mL of blocking buffer, containing 1% NDS (Sigma S30-100ML), 1% NGS (Sigma S26-100ML), 0.05% saponin (Sigma SAE0073), and 5% BSA (Sigma A7030-100G) in PBS, at room temperature for 1 hour. Cells were then incubated overnight at 4°C in 200 mL of blocking buffer containing primary antibodies. Cells were pelleted, washed with 200 mL of fresh PBS, before applying secondary antibodies diluted (1:1,000) in 150 mL of blocking buffer for 1 hour at room temperature. Cells were then re-pelleted, washed with 200 mL of PBS before resuspending in 300 to 400 mL of fresh PBS for flow cytometry. All post-fixation cell centrifugations were done at 650xg for 7 minutes at 4°C. A BD LSR Fortessa with HTS was used to quantify target fluorescence intensity per single living cell by gating for live cells, followed by selection for forward scatter (FSC) and side scatter (SSC) singlets. Cell gating and analysis was performed using FlowJo version 10.9.0.

### Membrane tension measurement using Flipper-TR

10,000 cells were plated in a μ-Dish 35 mm, high Glass Bottom microscopy dish (ibidi, 81158) before a 4 day serum withdrawal was started with media changes every other day. On day 4, cells were treated with 1 mM solution of Flipper-TR (Spirochrome CY-SC0202) in warmed Live Cell Imaging solution (ThermoFisher A59688DJ) for 15 minutes at 37°C. Fluorescence lifetime imaging microscopy (FLIM) was then done using a Leica SP8 Falcon FLIM confocal microscope with a HC PL APO CS2 63x/1.40 N.A. objective lens. A zoom factor of 1.81 was applied to get a pixel size of 100 x 100, a laser pulse setting of 80 MHz and a point scanning excitation speed of 600 Hz was used. Excitation was carried out at 488 nm, and emission was collected over a bandwidth of 570-650 nm using HyD SMD detector. For each experiment, 40 frames were acquired at a target rate of fewer than 1 photon per pulse to ensure adequate photon accumulation for lifetime analysis. A single z-plane with clear plasma membrane sections were selected for imaging^69^.

### Reactive astrocyte activation

2,000 to 5,000 cells were plated per well of a μ-plate 96-well square imaging plate (ibidi 89626-90) and a 4 day serum withdrawal was started with media changes every other day. On day 4, cells were treated with an activation cocktail of 3 ng/mL of IL1a (Peprotech AF-200-01A-100UG), 30 ng/mL of TNF-a (Peprotech AF-300-01a-250ug), and 400 ng/mL C1q (Complement Technology Inc A099) or vehicle of culture grade water and 0.1% BSA. After 24 hours of activation, standard 4% PFA fixation protocol was used along with standard the immunocytochemistry protocol for activation quantification.

### Lipid droplet staining and imaging

5,000 to 7,500 astrocytes were plated per well of a square 96-well imaging plate (ibidi 89626-90), in AM containing 2% FBS (ScienCell 1801). After 24 hours, the cell culture medium was changed to serum-free AM and the cells were cultured in this medium for 4 days with media changes every other day. Cells were then fixed using 4% paraformaldehyde (Electron Microscopy Sciences, EMT-5156) in DPBS for 15 minutes at room temperature. Fixed cells were incubated in PBS with phalloidin-647 and hoechst stain for 30 minutes at room temperature. Cells were then washed with PBS, 3 times for 15 minutes, before applying 100 μL of PBS with LipidTox Green for 30 minutes at room temperatures. Cells were directly imaged without washes using Nikon CSU-W1 Spinning Disk Microscope at 40X magnification, using NIS Elements version (Nikon).

### Fluorescence-activated cell sorting (FACS)

*APOE4* astrocytes were transduced with the aforementioned protocol and lentiviral constructs, using serum containing media for 72 hours. Cells were then cultured using fresh serum containing media and expanded to confluency in 2 x 10 cm tissue culture-treated dishes. When confluent, cells were collected using TrypLE Express (ThermoFisher Scientific, 12604039) and via FACS pure mCherry positive populations were isolated.

### Quantification and statistical analysis

Details of experiments, replicates, and data presentation can be found in figure legends. All experiments were performed in two human iPSC isogenic line sets and independent experimental conditions, transductions, and immune activations. To determine statistical significance, GraphPad Prism 10 software was used to perform two-tailed unpaired t-test, one-way ANOVA (with post hoc analysis to correct for multiple comparisons, details in appropriate legend), or Wilcoxon signed-rank test. Data are represented as box-and-whisker graph with outer whiskers representing minimum to maximum, box edges representing first and second quartiles, median represented by a line, and + denoting mean. Individual data points for each graph were also superimposed, details in each figure legend. FlowJo (version 10.9.0) was used to process fluorescence-activated cell sorting data. Imaging processing was performed using Nikon NIS-Elements AR software version 6.10.01.

### Analysis of flow cytometry

FlowJo version 10.9.0 was used to gate cells and quantify the proportion of HA-positive cells, mean intensity of HA expression, mean intensity of fluorescently conjugated transferrin ligand, mean intensity of EEA1 (early endosome) intensity. Mean values from experimental conditions (*APOE4* astrocytes, *APOE4* astrocytes transduced with adaptor proteins) were normalized to mean values for control conditions (*APOE3* astrocytes, *APOE4* astrocytes transduced with dark GFP). Details for normalizations are in figure legends.

### Analysis of immunocytochemistry

All immunocytochemistry image acquisition was done with a Nikon CSU-W1 Spinning Disk Microscope and all analysis was using Nikon NIS-Elements AR software. For intensity measurements, cells from a replicate were imaged using the same laser conditions (percent power, exposure time) and imaged on the same day. Nikon NIS-Elements AR software was used for semi-automated analysis of confocal images. A maximum intensity projection was generated per frame and thresholding was applied to removed background noise. Nuclear (Hoechst) and whole cell (vimentin, phalloidin) stains were used to mask the appropriate regions of interest for intensity quantification and normalization of intensity measurements to cell area per frame. Sum intensities per area were normalized to average sum intensity per area of control conditions, details in figure legends.

### Analysis of plasma membrane tension using Flipper-TR

Flipper-TR membrane tension analysis was done using Leica LAS X Life Science Microscope Software Platform (Version 3.5.7.2), where a threshold was applied to remove background signal and regions of interest were drawn around bright clear plasma membranes of cells. Fluorescent lifetime (ns) for regions of interest were quantified using phasor analysis^70^ and all experimental values were normalized to average of control (*APOE3* or *APOE4* transduced with darkGFP), details in figure legends.

### Analysis of Neurolipid Atlas whole cell lipidomics

Whole cell lipidomics data for fatty acid chain length (number of carbons) and unsaturation (number of C=C double bonds) was downloaded from the NeuroLipid Atlas website. Number of carbons and number of C=C double bonds for every PA, PS, PI, PG, PC, PE, and SM from 3 replicates and from 3 lines (two KOLF2.1J clones and BIONi-037A) were plotted together in a volcano plot. Negative log 10 of the adjusted p-value per lipid species was plotted on the y-axis and the log2 fold change (*APOE3* vs *APOE4*) were plotted on the x-axis.

### Analysis of Raman spectroscopy

Samples used for Raman spectroscopy were transferred into μ-Dish 35 mm, high Glass Bottom (ibidi) and cultured (as described above), fixed with 4% paraformaldehyde in dPBS, and sealed with a round glass coverslip. Raman spectra were acquired using a DXR2xi Raman microscope (ThermoFisher Scientific) with 10 mW of 532 nm laser directed at the sample through a 100X oil confocal objective (N.A. = 1.3 a working distance of 0.2 mm UPLanFLN Olympus America Inc.) at 2 s exposure time, through a 50 μm pinhole, for a pixel size between 200 nm and 1 μm, for the 100 to 3400 cm-1 spectral region. Chemical images were produced by the peak height selection of the assigned peaks using Thermo Fisher Scientific OMNICxi software (ThermoFisher Scientific), using a local selected baseline correction and displayed as a heatmap. Spectra from representative random regions of the cellular membrane were selected for spectral extraction. The edges of the spectra between 200 – 500 cm-1 and 3100 – 3400 cm-1 were eliminated, as they do not contain any relevant biological information. Next, an automated background subtraction function was applied, followed by normalization to the sum and finally spectra were overlayed using OriginPro 2021b (Origin Lab Corporation). An average spectrum was highlighted from at least five spectral regions of the membrane. The regions of interest were plotted and saved as excel files and replotted in Graph Pad Prism (GraphPad Software, LLC) for further analysis and display.

### Analysis of platinum replicas

Montages were stitched together using IMOD. File names were blinded so that subsequent quantification could be done without bias. Clathrin in the stitched images was automatically segmented. Structures were binned into classes by degree of curvature (flat, dome, or sphere). These automatic functions were performed using deep learning model Mask R-CNN and image analysis platform Napari using the previously published method^71^. Automatic segmentation was then manually verified and corrected using Napari. Cell masks were also generated using Napari. Clathrin structure class compositions graphs and statistical analysis was done using Prism. Prism ROUT method, with a Q=0.1 %, was used to identify and remove outliers.

## Author Contributions

Conceptualization, A.S.G., L.G., J.T., and P.S.N.; Methodology, A.S.G., L.G., C.C., A.L., C.C.L., M.L., J.T., and P.S.N.; Software, A.S.G., L.G., C.C.; Validation, A.S.G., L.G., A.L.; Formal Analysis, A.S.G., L.G., C.C., A.L.; Investigation, A.S.G., L.G., C.C., A.L.; Data Curation, A.S.G., L.G., L.G.Y., and P.S.N.; Writing – Original Draft, A.S.G., L.G., and P.S.N.; Writing – Review and Editing: all authors, Resources, C.C., A.L., L.G.Y., J.T.R., M.L., J.T., and P.S.N.; Supervision, C.C., L.G.Y., M.L., J.T., and P.S.N.; Funding Acquisition, M.L., J.T., and P.S.N.

The contributions of the NIH authors were made as part of their official duties as NIH federal employees, are in compliance with agency policy requirements, and are considered Works of the United States Government. However, the findings and conclusions presented in this paper are those of the authors and do not necessarily reflect the views of the NIH or the U.S. Department of Health and Human Services.

## Acknowledgement

We thank members of the Narayan Lab, as well as Drs. Richard Proia, Ashley Frakes, Richard Youle, Elise Marsan, Anne Hart, Michael Ward, and Mark Cookson for helpful discussions of this work. We also thank the NHBLI Flow Cytometry Core for their help with many elements of this project. Figures 1 and 4 were created with BioRender. This research was supported by the Intramural Research Programs of the National Institute of Diabetes and Digestive and Kidney Diseases (NIDDK) (1ZIADK075158), the National Heart Lung and Blood Institute (NHLBI), and the National Cancer Institute (NCI) within the National Institutes of Health (NIH).

---

**Supplemental Figure 1:**
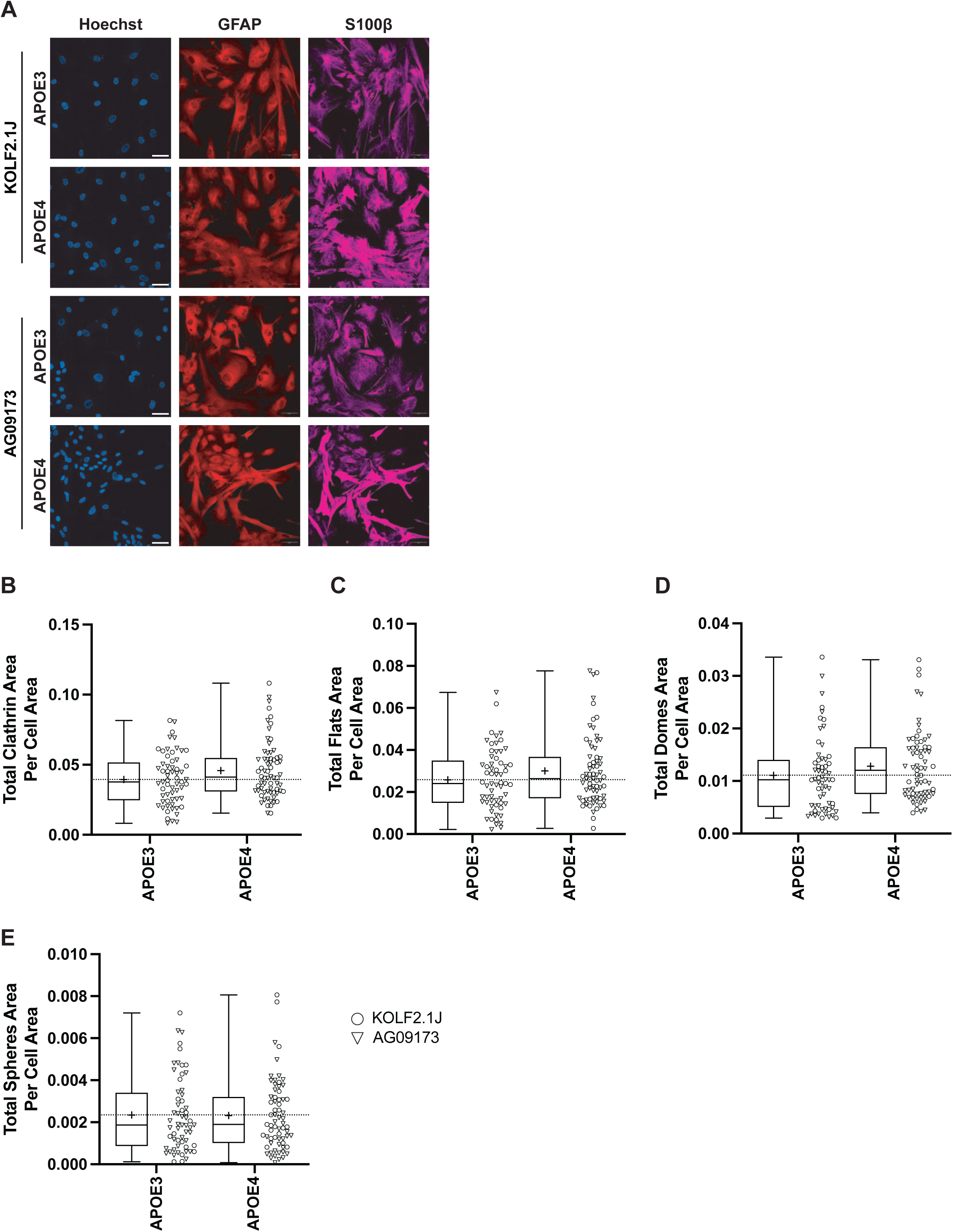
iPSC line quality control and further PREM characterization. **A)** GFAP and S100β immunocytochemistry for KOLF2.1J and AG09173 iPSC-derived astrocytes (*APOE3* and *APOE4* homozygous lines). Scale bar is 50 µm. Quantification of **B)** total clathrin area, **C)** total flat area, **D)** total dome area, and **E)** total sphere area as a function of cell membrane area in membranes imaged with PREM. N=61-68 cells per genotype, across two iPSC lines (KOLF2.1 O, AG09173 ∇). Box and whisker, minimum to maximum with line at median, cross (+) denotes mean alongside individual cell data. Unpaired t-test used for statistical analysis.

**Supplemental Figure 2:**
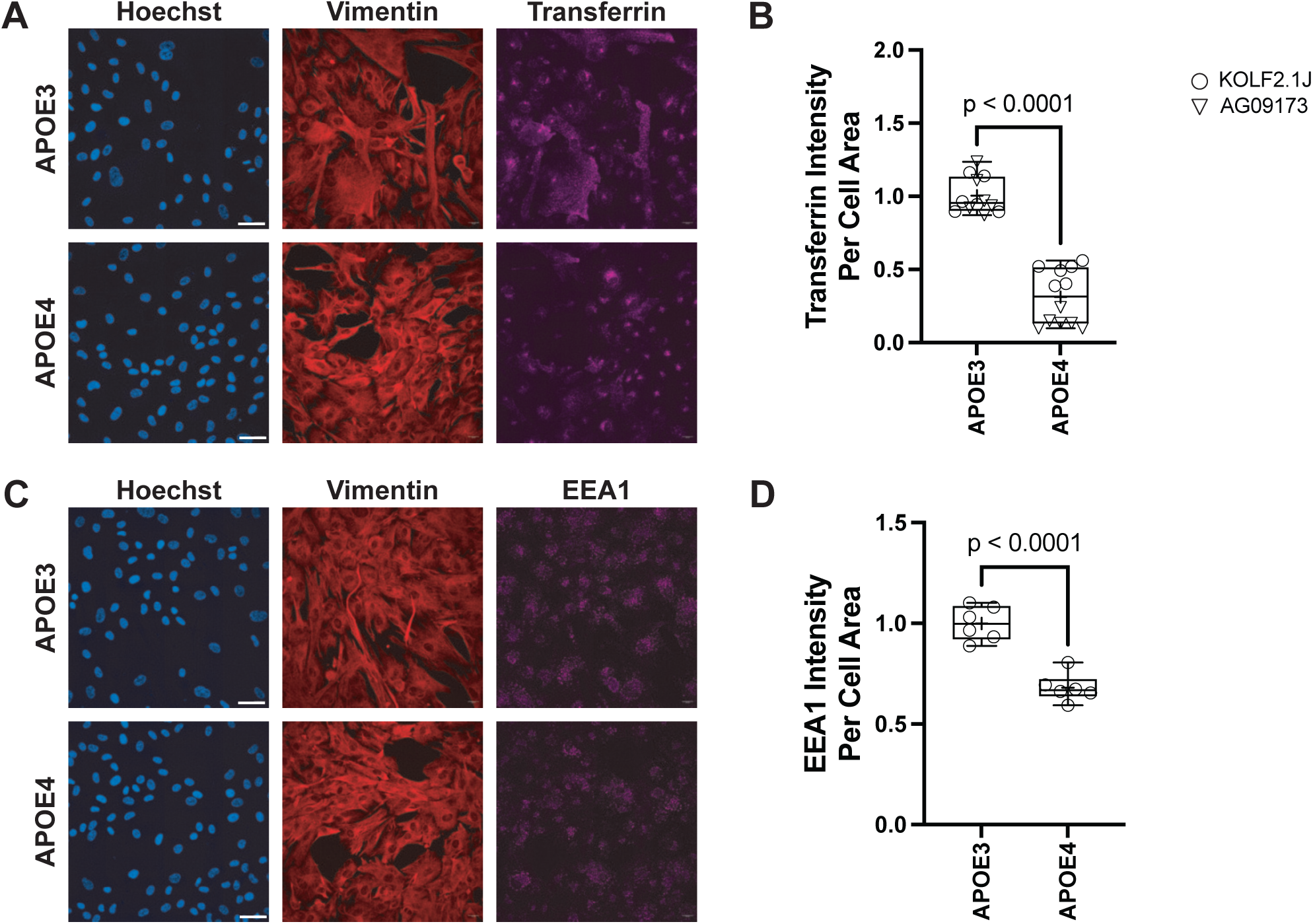
Immunofluorescence images of transferrin uptake and EEA1 endosomes in iPSC-derived astrocytes. Representative images for **A)** cellular uptake of fluorescent transferrin with **B)** quantification of fluorescent intensity per cell, N=12 wells per genotype, across two iPSC isogenic lines (KOLF2.1 O, AG09173 ∇). Representative images for **C)** EEA1 positive early endosomes with **D)** quantification (N=6 wells per genotype for KOLF2.1J). Box and whisker, minimum to maximum with line at median, cross (+) indicates mean. Unpaired t-test used for statistical analysis. All scale bars are 50 µm.

**Supplemental Figure 3:**
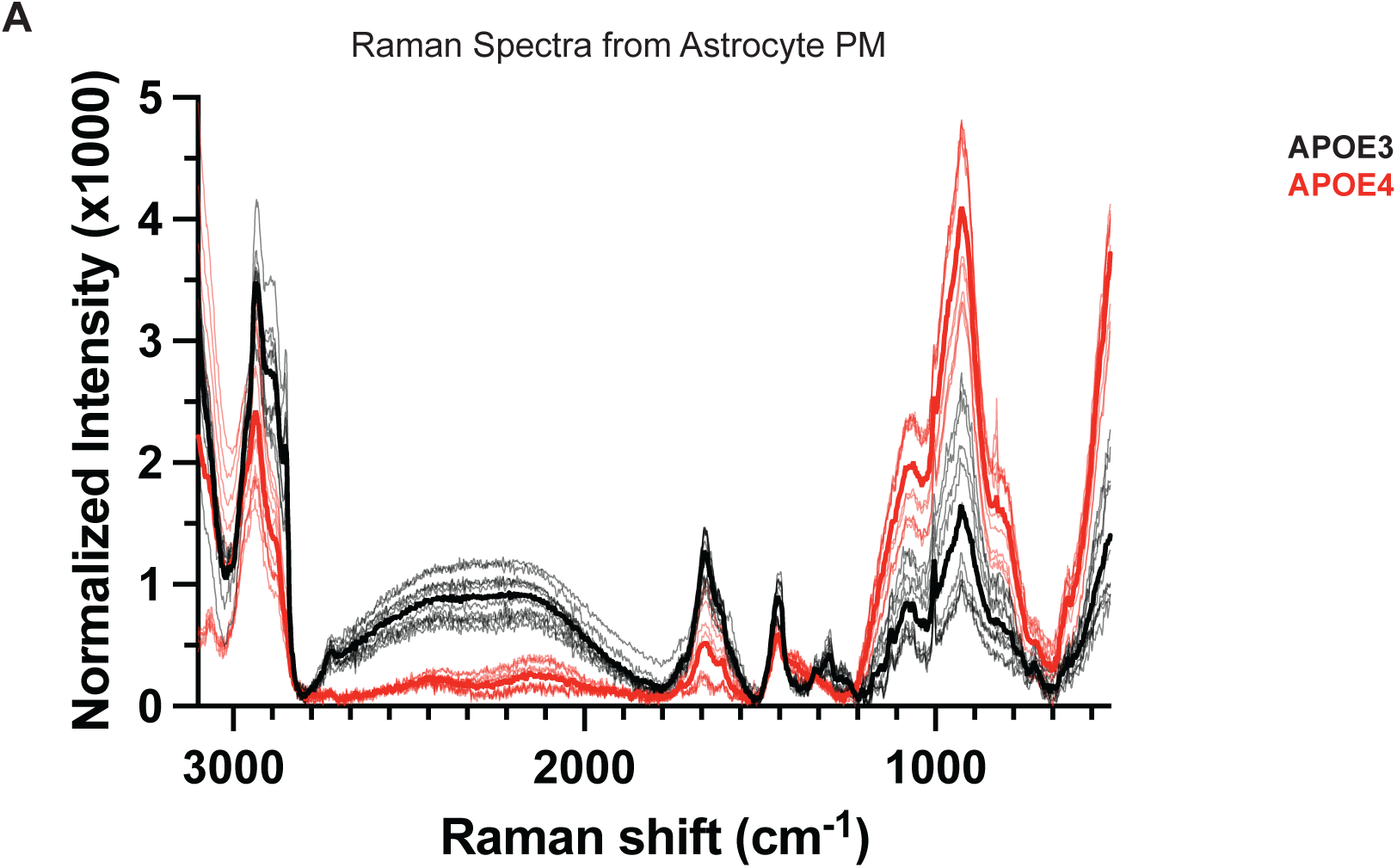
Complete Raman spectra from *APOE3* and *APOE4* astrocyte plasma membrane regions. **A)** Full Raman spectrum from 500 cm^-1^ to 3100 cm^-1^, from N=10 membrane regions from *APOE3* astrocytes (grey, solid black = *APOE3* average) and from N=10 membrane regions from *APOE4* astrocytes (pink, solid red = *APOE4* average)

**Supplemental Figure 4:**
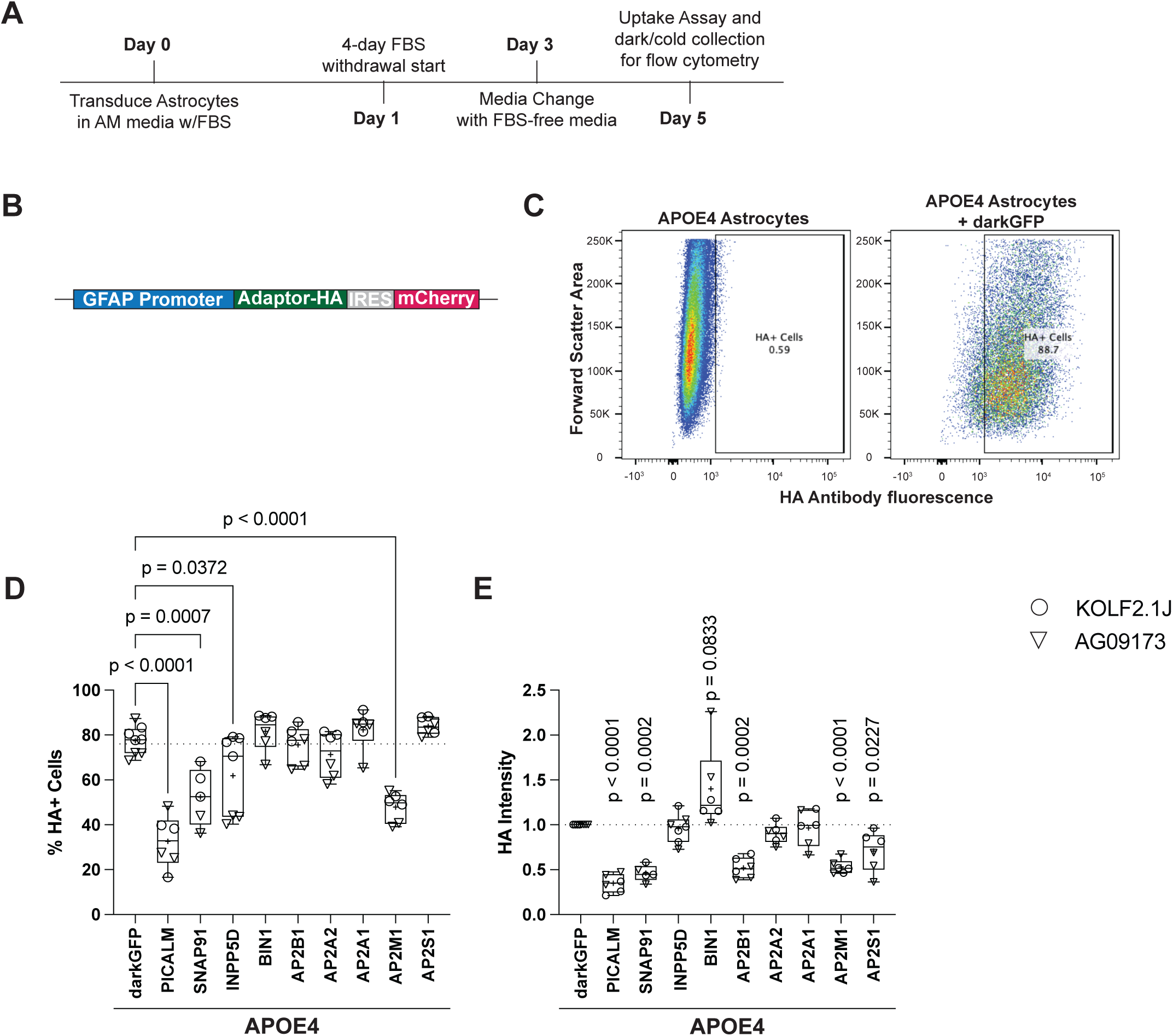
Exogenously expressed protein expression levels. **A)** Experimental workflow for *APOE4* astrocyte lentiviral transduction and culture in serum free media before endocytosis quantification via flow cytometry. **B)** Diagram of lentiviral construct used to transduce and express each candidate gene of interest (GOI) tagged with HA in *APOE4* astrocytes. **C)** Gating example to identify positively transduced *APOE4* astrocytes based on HA positivity. Box and whisker plot (minimum to maximum values) for quantification of multiple flow cytometry runs for **D)** percentage of transduced cells (quantified by number of singlet cells that are HA positive), N=5-7 flow cytometry runs per endocytosis protein, across two iPSC lines (KOLF2.1J, AG09173), one-way ANOVA with multiple comparisons used for statistical analysis. **E)** HA tag intensity from positively transduced astrocytes, N=5-7 independent flow cytometry runs per endocytosis protein, across two iPSC lines (KOLF2.1 O, AG09173 ∇) flow cytometry means normalized to *APOE4* astrocytes transduced with control *darkGFP* values from same run, Wilcoxon signed-rank test.

**Supplemental Figure 5:**
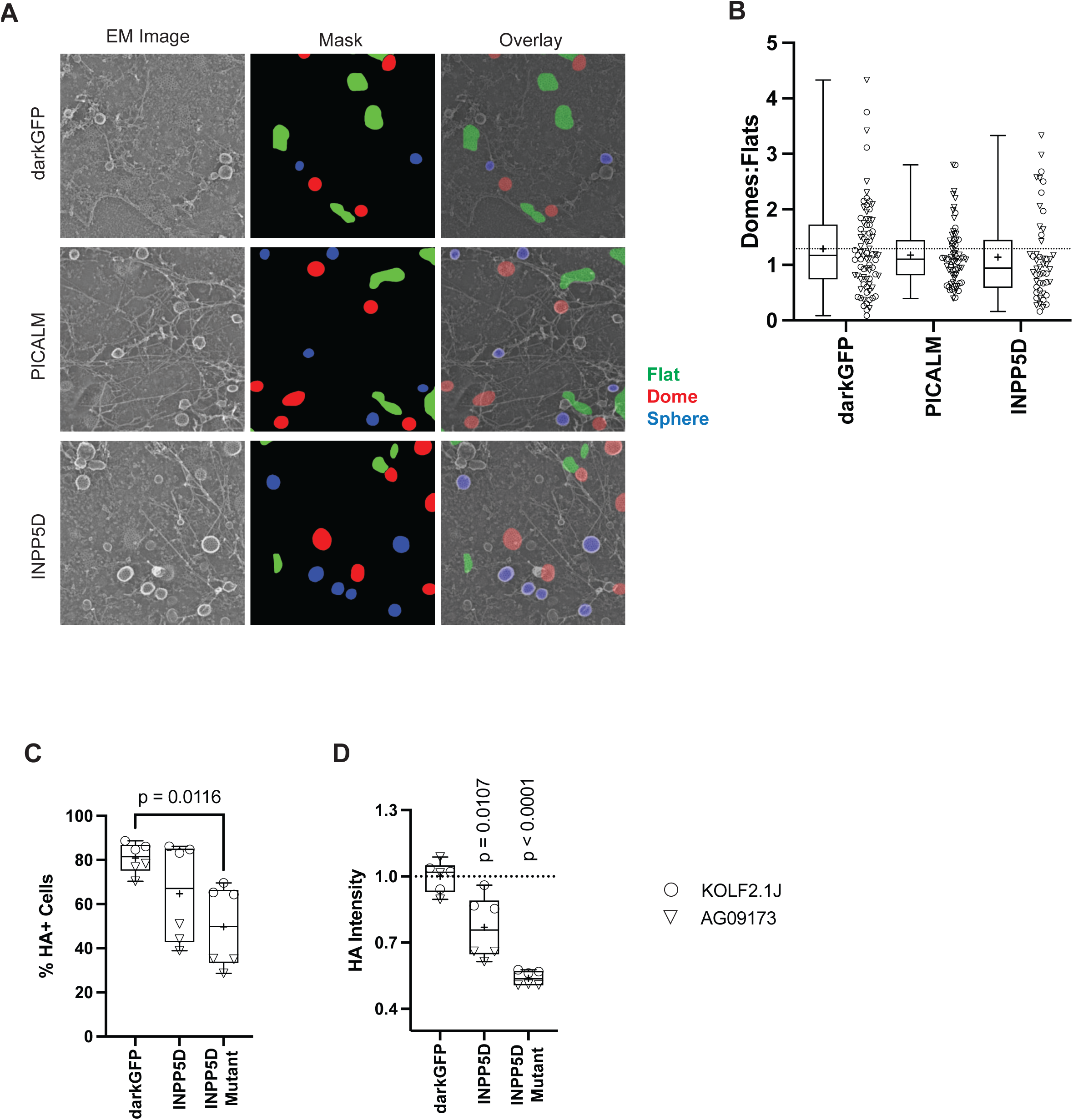
Adaptor PREM and mutant expression levels. **A)** Example images of PREM from *darkGFP*, *PICALM*, or *INPP5DI*, with flat, dome, and spherical clathrin structure masks used for quantifications. **B)** PREM quantification of number of Domes:Flats ratio in *darkGFP*, *PICALM*, or *INPP5D* expressing *APOE4* astrocytes. N=44-81 cells per condition, across two iPSC isogenic lines (KOLF2.1 O, AG09173 ∇). Box and whisker, minimum to maximum with line at median, cross (+) indicates mean. One-way ANOVA with multiple comparisons used for statistical analysis. **C)** Box and whisker plot (minimum to maximum values with line at median) for quantification of multiple flow cytometry runs for percentage of transduced cells (quantified by number of singlet cells that are HA positive), N=6 flow cytometry runs per condition, across two iPSC lines (KOLF2.1J, AG09173), one-way ANOVA with multiple comparisons used for statistical analysis. **D)** HA tag immunofluorescence intensity from positively transduced astrocytes, N=6 flow cytometry runs per condition, across two iPSC lines (KOLF2.1 O, AG09173 ∇) flow cytometry means normalized to *APOE4* astrocytes transduced with control *darkGFP* values from same run, Wilcoxon signed-rank test.

## Notes

### Competing Interest Statement

The authors have declared no competing interest.

